# Polarization light-sheet microscopy and tomography (PLµTo) for flow-based imaging of 3D microcarrier mesenchymal stem cell culture

**DOI:** 10.1101/2025.01.02.631120

**Authors:** Oscar R. Benavides, Berkley P. White, Roland Kaunas, Carl A. Gregory, Alex J. Walsh

**Affiliations:** Department of Biomedical Engineering, Texas A&M University, College Station, Texas, United States of America; School of Medicine, Texas A&M Health Science Center, Bryan, Texas, United States of America

**Keywords:** Optical microscopy, light-sheet, elastic scattering, imaging cytometry, bioreactor-microcarrier cell culture

## Abstract

Light-sheet microscopy is an increasingly popular imaging tool for studying biological processes, however, it generally requires the use of exogenous fluorescent contrast agents which can be toxic to live cells. Light-sheet tomography is the label-free analog of light-sheet microscopy, utilizing elastic scattering contrast to visualize the structure of the biological sample. Here, we present a multi-modal polarization light-sheet microscopy and tomography (PLµTo) system for on-line, non-destructive, and non-invasive volumetric imaging of human mesenchymal stem cells, also referred to as mesenchymal stromal cells, attached to spherical gelatin methacryloyl microcarriers for image analysis-based quantitative monitoring of adherent stem cell culture expansion. The PLµTo system, which is built on an inverted selective plane illumination configuration, is compatible with standard horizontal sample mounting and microfluidics for multimodal volumetric data acquisition. This work is the first demonstration of scattering-based volume optical imaging flow cytometry.

## Introduction

Single plane illumination microscopy (SPIM), or light-sheet microscopy (LSM), is a fast and photo-efficient wide-field technique for high spatial and temporal resolution imaging of biological samples. High resolution three-dimensional (3D) reconstructions of the sample can be generated as this method achieves optical sectioning due to the thin plane of illumination within the sample which minimizes the contributions of out-of-focus light to image formation. Light-sheet fluorescence microscopy (LSFM) utilizes the generation of fluorescence contrast and is increasingly popular for studying the development of multicellular organisms due to the imaging speed, spatial resolution, and low-light exposure advantages of light-sheet microscopy [1], [2], [3], [4]. Fluorescence-based assays, however, generally require the use of exogenous fluorescent markers, which can be toxic to cells and incompatible with live cell or organism studies, to generate biological signals of interest. Fortunately, light-sheet tomography (LST) presents a label-free alternative to LSFM where contrast is generated by detecting elastically scattered photons from the illuminated sample [5–10]. This imaging technique has been used to study the development of plant roots [5], the structure of rodent brains [6], and more recently to visualize and quantify proliferation of 3D bioreactor-microcarrier mesenchymal (stem/stromal) cell (MSC) cultures [9], [11]. As a label-free imaging technique, LST is non-invasive, non-destructive, and compatible with live cell imaging, making it an attractive solution for monitoring of bioreactor-microcarrier cell cultures.

Mesenchymal stem cells (also referred to as mesenchymal stromal cells) are attractive cytotherapeutic candidates, based on their well-documented anti-inflammatory and immunomodulatory capabilities [12–14]. One significant challenge in the manufacturing of cytotherapies is the tremendous quantities of cells needed for multiple patient doses in clinical trials. This has spurred the development of novel large-scale suspension culture methods that can efficiently generate high cell yields. relative to 2D monolayer cultures without compromising viability, identity, or differentiation potential [15].

Unfortunately, the visualization-based assays that are standards for monolayer cultures do not readily translate to 3D microcarrier-based cultures. The development of novel cell expansion strategies simultaneously motivates the advancement of analytical tools to study these 3D cell cultures. In fact, the Food and Drug Administration (FDA) has encouraged pharmaceutical manufacturers to develop and deploy Process Analytical Technologies (PATs) for near real-time monitoring of key variables throughout the manufacturing process to ensure safety, efficacy, and quality of the final product [16]. An ideal PAT system for microcarrier cultures would perform 3D imaging of cells encompassing the entire microcarrier surface in near real-time using non-destructive label-free contrast to assess the health of the culture through visualization of cell morphology and enumeration of cells to track proliferation. Our previous work has demonstrated that MSCs cultured on hydrogel microcarriers can be visualized and quantitatively characterized using light-sheet tomography (LST) for non-destructive, non-invasive, and label-free elastic scattering contrast imaging [9].

A closed-loop sampling system is necessary for PATs to maintain the sterility and yield of the bioreactor culture while monitoring the proliferation of MSCs. Here, on-line flow-based monitoring was demonstrated by imaging MSCs cultured on hydrogel microcarriers in a 10 mL rotating wall vessel (RWV) bioreactor through FEP tubing and a microfluidic device with an FEP cover. The research reported here presents the construction, characterization, optimization, and application of a custom-built polarized light-sheet microscopy and tomography (PLµTo) system that enables on-line, non-destructive, and non-invasive volumetric imaging of cells attached to microcarriers for image analysis-based quantitative monitoring of MSC expansion. Image quality can be improved by controlling the polarization states of the illumination and detection paths to minimize detection of FEP-scattered photons and maximize MSC SNR [7], [8], [17]. The optical performance of the system was characterized and optimized using fluorescent spheres and fixed hydrogel microcarrier-cell samples. The PLµTo system, which is built on an inverted selective plane illumination configuration, is compatible with standard horizontal sample mounting and hydraulic flow for multimodal volumetric data acquisition.

## Methods

### PLµTo system design

The system was designed to permit on-line, flow-compatible, sub-cellular resolution fluorescence and tomographic imaging of cells on microcarriers using fluorescence and elastic scattering contrast. Fluorescence capabilities were included with the elastic scattering contrast to enable a direct comparison of scattering contrast images and data with gold-standard fluorescence-based visualization of cell morphology. A schematic of the system is presented in Figure 1. A 488 nm laser diode (Coherent OBIS 488 nm 40 mW) and 633 nm laser diode (Coherent OBIS 633 nm 70 mW) are used for illumination sources. The two beams are steered to be concentric after t the dichroic mirror (DM) (Thorlabs DMLP505) and then magnified 2x by passing through 25 (Thorlabs AC254-025-A) and 50 (Thorlabs AC254-050-a) mm focal length lenses. A λ/2 plate (Newport 10RP42-1) or λ/4 plate (Edmund Optics 390-32) is placed in the optical path to create linear vertical, linear horizontal, right circular, or left circular illumination polarization states. A Powell lens (Thorlabs LGL175) is used to generate a laser line with a flat-top, as opposed to gaussian, intensity profile [18]. Two cylindrical lenses (Thorlabs ACY254-050-A) collimate the fast axis and focus the slow axis of the beam, which is imaged onto the back focal plane of the 10X 0.3W NA illumination objective (Olympus UMPLFLN 10X) that is oriented at 45° with respect to the horizontal plane. A variable vertical slit (S) placed at the common focal plane of the cylindrical lenses is used to control the illumination NA. The emitted signal is collected by the 20X 0.5W NA detection objective (Olympus UMPLFLN 20X) that is orthogonal to the light-sheet plane. An emission filter to block out 488 nm illumination (Semrock BLP01-488R-25) or 633 nm illumination for fluorescence imaging (Semrock LP02-638RU-25) is placed in the objective infinity space. A second linear polarizer (Thorlabs WP50L-UB) is placed in infinity space to change the detected polarization states. A 150 mm tube lens (Thorlabs AC254-150-a) images the field of view onto the camera (pco.edge 5.5)

**Figure 1.**
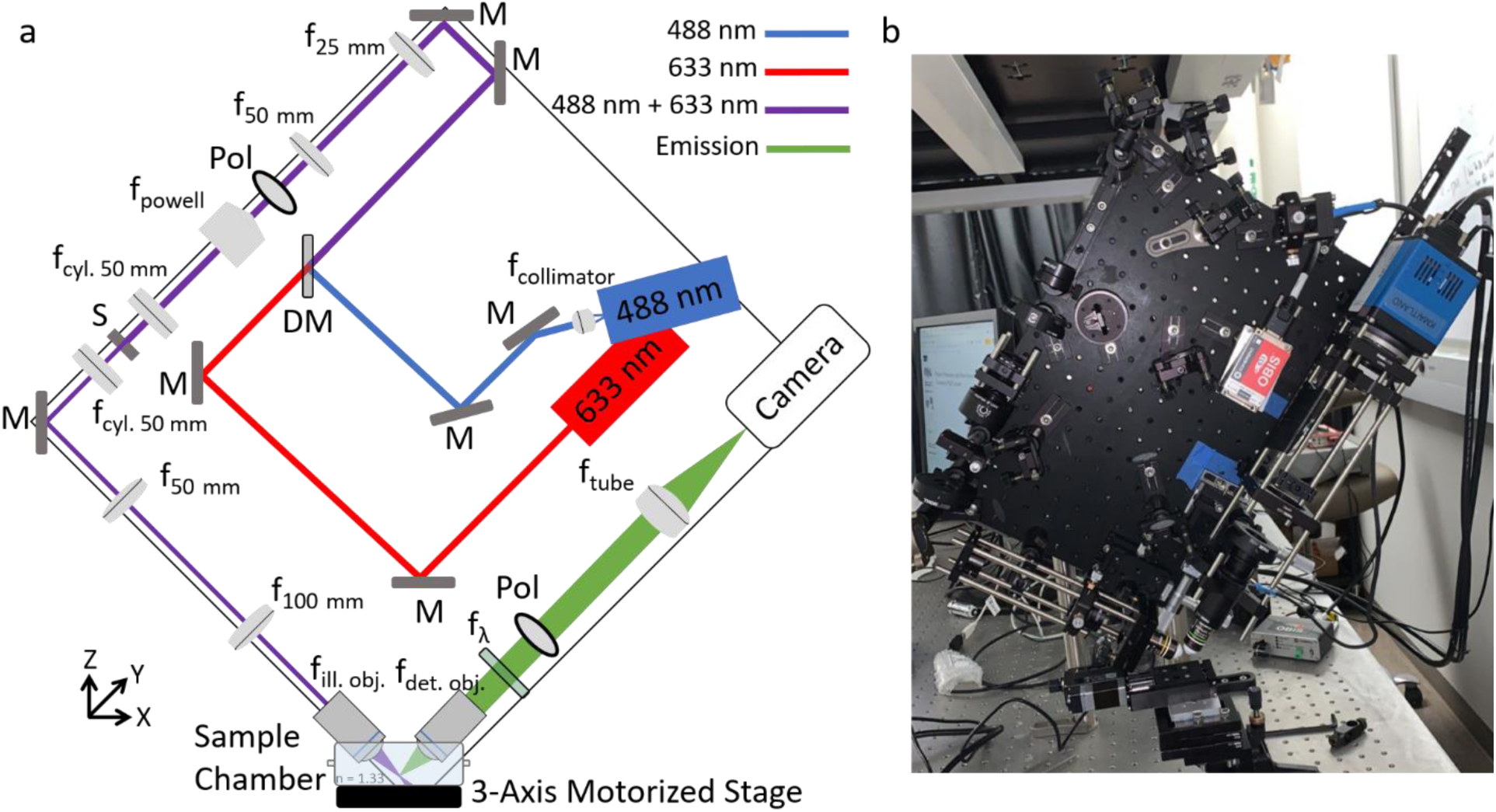
a) Schematic of the custom-built PLµTo system. f = lens, M = mirror, Pol = polarizer, S = slit, DM = dichroic mirror. b) Photograph of PLµTo system.

### Data acquisition and processing

The default camera software, pco.camware (PCO v4.14), was used to collect data. Owing to the oblique imaging geometry in relation to the sample motion, the acquired dataset is skewed if the oblique angle is not accounted for (Figure 2a). Image visualization software incorrectly displays the data without properly pre-processing the data (Figure 2b,c). De-skewing of the volumetric data can be performed by linearly displacing pixels a pre-set number of frames determined by the illumination angle and axial step size, or by an affine transformation to scale and shear the original voxels to the correct dimensions and locations (Figure 2d) [19], [20]. Here, the de-skewing was done post-acquisition using ImageJ software [21] and the TransformJ plugin [22] to perform the affine transformation (Figure 2e-g).

**Figure 2.**
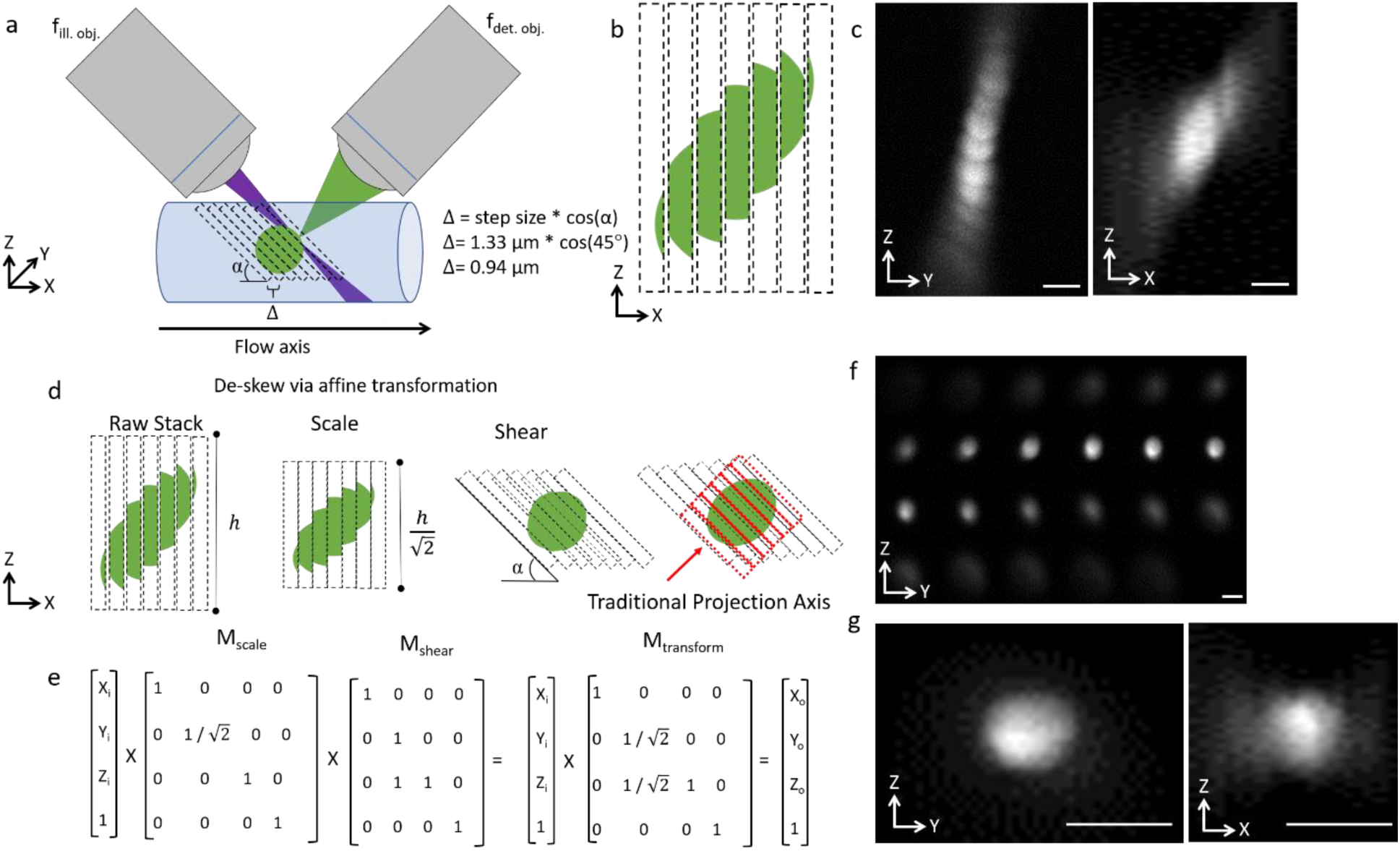
Illustration of the de-skewing method used for correctly reconstructing the 3D imaging data. a) Oblique image acquisition requires de-skewing for accurate image analysis. B) The de-skewed stack will appear distorted when visualized. c) Maximum intensity projections of 10 μm fluorescent beads without de-skewing. d) De-skewing is achieved using an affine transformation that scales and shears the volume. A traditional en-face projection can be acquired by rotating the object. e) The matrix math performed in the affine transformation. f) A montage of slices along the red ROI in d showing a traditional en face view of the sphere. g) Maximum intensity projections of the traditional en face view Scale bar = 10 μm.

### Optical characterization

The optical performance of the system was characterized by measuring the point-spread function (PSF) using 100-nm multi-spectral fluorescence beads (ThermoFisher TetraSpeck Microspheres T7279) and ImageJ software. A 1 µL solution of 0.1 µm fluorescent beads were suspended in 5 mL of 1% agarose and then deposited into a 3D-printed sample mount [23]. The agarose embedded samples were imaged using 488 nm illumination (20 mW), 29 µm/s stage scan, and 75 frames per second (FPS) acquisition rate. The volumes were de-skewed, and maximum intensity projections were created (Figure 3b). The full width at half maximums (FWHM) of the line profiles through the maximum intensity projections were fit with a one-dimensional Gaussian function, and were characterized at various depths within the agarose sample (Figure 3c).

**Figure 3.**
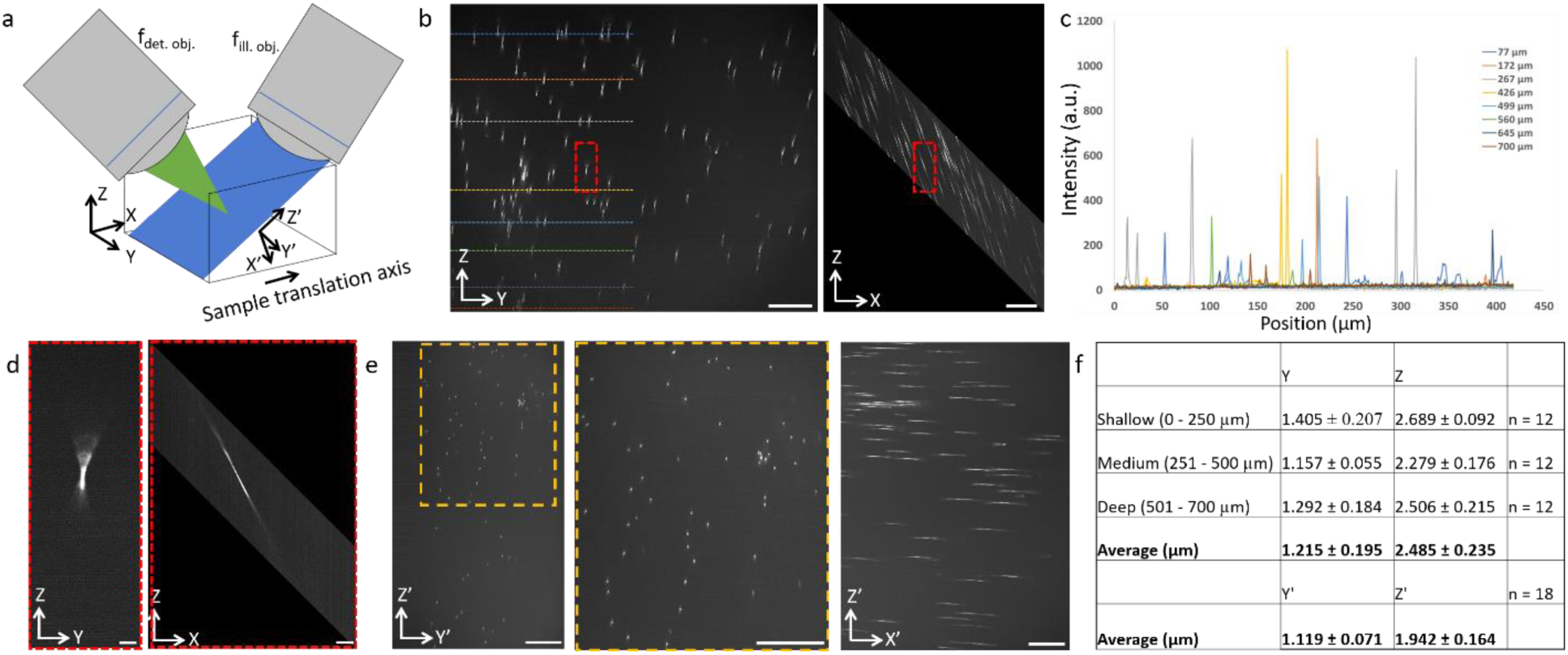
(a) Illustration of the spatial coordinates and the rotated coordinates along the acquisition angle. b) Maximum intensity projections of 0.1 μm beads. The full field of view is approximately 700 x 840 μm^2^ (depth x width). C) The intensity of beads is characterized up to a depth of 700 µm in the agarose-embedded sample (dashed lines). Scale bar = 100 μm. d) A zoomed-in ROI from b) showing a single bead (red ROI). Scale bar 10 μm. e) Maximum intensity projections along the oblique angle showing a traditional en face view, zoomed in ROI, and the orthogonal projection. Scale bar = 100 μm. f) Measured spatial resolution along varying depths through the sample, and measured along the oblique acquisition angle projection.

### Induced pluripotent stem cell-derived hMSC (ihMSC) culture

The ihMSCs were cultured in a 10 mL rotating wall vessel (RWV) bioreactor (RCCS-8DQ bioreactor, Synthecon) on custom-fabricated 120 ± 6.2 μm diameter gelatin methacryloyl (gelMA) microcarriers, as previously described [9]. Briefly, passage 4 ihMSCs were expanded in monolayer cell culture to obtain the required cell numbers. gelMA microcarriers with a combined growth area of 50 cm^2^ (∼110,000 microcarriers) and 5×10^4^ cells (1000 cells/cm^2^) were incubated in 10 mL of complete culture medium (CCM) (α-Minimum Essential Medium, 10% fetal bovine serum, 2mM L-glutamine, 100 U/mL penicillin, and 100 μg/mL streptomycin) in the RWV bioreactor at 24 revolutions per minute. Half of the media was replaced with fresh CCM every 2 days.

### Polarization optimization

By using linear polarizers, a λ/4 waveplate, and a λ/2 waveplate, 12 different system polarization state configurations were evaluated to optimize the elastic scatting image contrast. The illumination polarization states were linear vertical, linear horizontal, right circular, and left circular; the detection polarization (analyzer) state was either no polarization, linear vertical, or linear horizontal. The SNR for the different system polarization states was measured from single images of passage 4 day 3 ihMSCs on gelMA microcarriers embedded in agarose within FEP tubing and the FEP-covered chip. Cell SNR was measured as the ratio of cell cytoplasm intensity to the background agarose intensity. The FEP scattering was characterized as the ratio of mean scattering intensity from the FEP tubing or sheet compared to the surrounding agarose or de-ionized water (DI) water inside the tubing [7], [8]. The speckle contrast (S.C.) of the FEP tubing was also characterized as the ratio of the intensity standard deviation compared to the mean intensity.

### Stage scanning and flow-enabled imaging

To evaluate the quality of images that PLµTo acquires of complex cell cultures, ihMSCs attached to gelMA microcarriers were immobilized in agarose in FEP tubing and imaged using stage scanning to simulate flow translation. Cell number and cell volume were quantified using previously published Imaris-based image analysis methods [9]. To demonstrate the applicability of PLµTo as an on-line PAT for monitoring microcarrier-based cell cultures, a syringe pump (Legato 180, KD Scientific) and FEP tubing (McMaster Carr 8703K114) were used to flow formalin-fixed microcarrier-cell samples at 1,000 nL/min through the PLµTo system.

## Results

### PSF characterization

The oblique acquisition causes the ZY and ZX projections of the de-skewed volume to be reconstructed at the acquisition angle (Figure 3d). However, a more intuitive view can be seen by taking a projection along the Z’Y’ and Z’X’ axes, which gives a view of the sample similar to standard trans- and epi-illumination microscopes (Figure 3e). The average lateral resolutions across the entire FOV were 1.215 ± 0.195 and 2.485 ± 0.235 along the ZYX spatial coordinates, and 1.119 ± 0.071 and 1.942 ± 0.164 (mean ± standard deviation) along the rotated Z’Y’X’ coordinates (Figure 3f).

### FOV characterization

The detection arm of the system is oriented at 45° with respect to the collection, or axial, axis and is comprised of a 20X 0.5W NA objective lens and 150 mm focal length lens which produce an effective magnification of 16.667X. The sCMOS camera has a 2560 x 2160 pixel array and pixel size of 6.5 x 6.5 µm^2^, which correlates to a de-skewed lateral FOV of ∼840 x 700 µm^2^ (w x h) (Figure 3b). Fluorescence signal from 100 nm spheres can be resolved at depths through 700 µm (Figure 3c).

### Polarization filtering

The polarization states of the illumination and detection arms were varied to optimize the label-free imaging protocol for passage 4 day 3 iH-MSCs attached to gelMA microcarriers through a 200 µm thick FEP sheet-covered microfluidic chip. All images within Figure 4 subfigures (a, b, and c) are displayed at the same gray-scale color map and contrast level. Left circular polarized illumination and vertical polarized detection provided the greatest SNR of 12.61 (Figure 4a). The linear vertical polarized illumination and co-polarized detection state similarly provided high cell SNR. The use of linear horizontal polarized state for either illumination or detection paths resulted in poor SNR. Linear vertical illumination states minimized FEP scattering compared to circular illumination configurations, and the linear horizontal detection configuration minimized the collection of FEP- and cell-scattered photons (Figure 4b). The polarization-dependent scattering of 1 mm I.D. FEP tubing with 250 µm thick walls was also characterized (Figure 4c). The linear horizontal detection configuration permitted visualization of ballistic photons scattered at the FEP-water boundary. Linear vertical detection states produced less scattering from the FEP tube than linear horizontal detection. The speckle contrast within the FEP tube was minimized using linear vertical detection. Linear vertical and left circular illumination states caused greater FEP scattering than linear horizontal or right circular illumination states.

**Figure 4.**
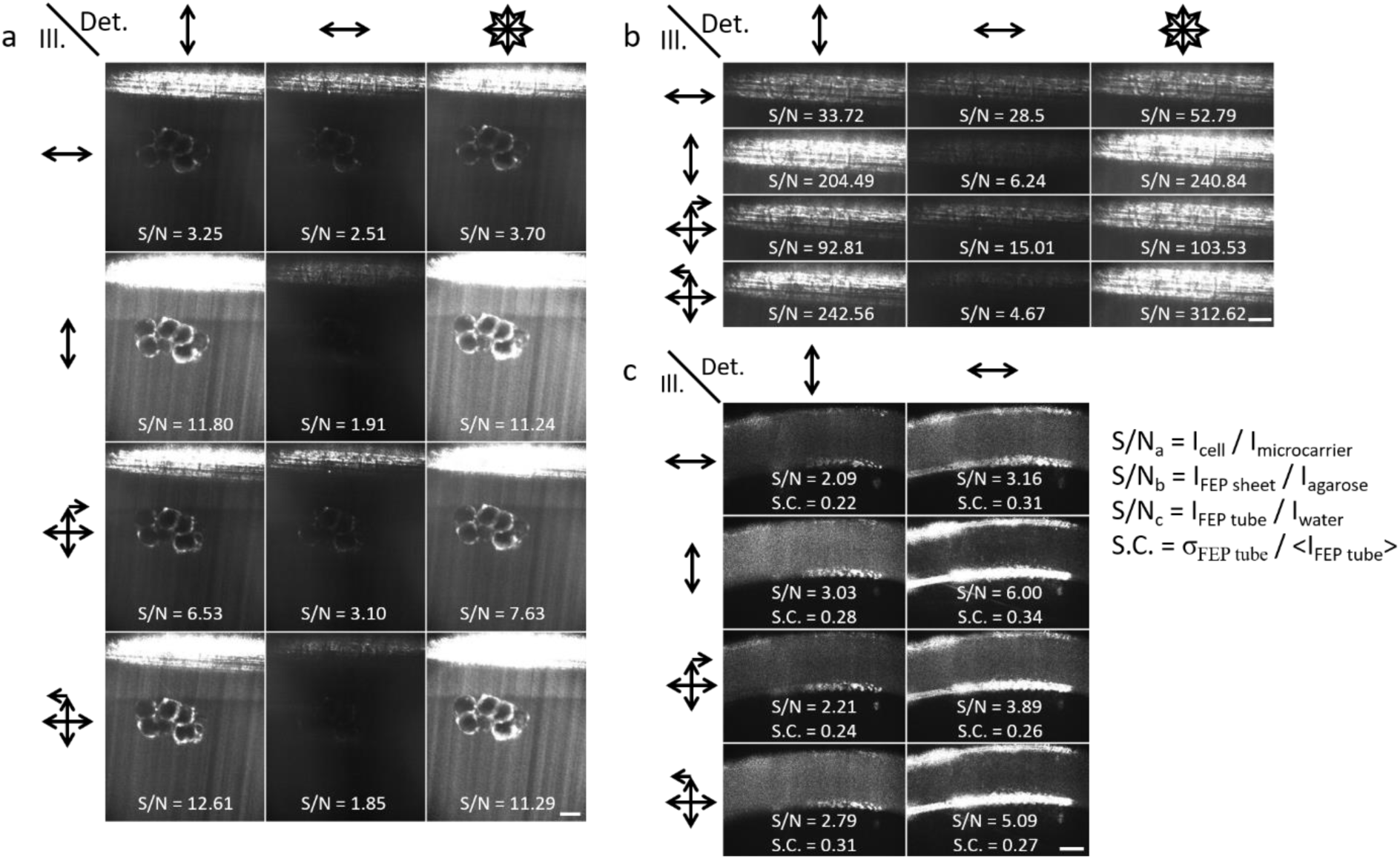
Passage 4 day 3 iH-MSCs attached to gelMA microcarriers imaged using 633 nm elastic scattering and various polarization configurations in the illumination and detection arms. The overlay values represent SNR. a) Co-polarization with vertical polarization produced the greatest SNR. b) Copped regions of interest showing the effects of polarization on the FEP sheet. Contrast of images in b rescaled from a but consistent within b for visualization of FEP tube scattering. (c) Polarization filtering test on 1 mm FEP tubing submerged and filled with de-ionized (DI) water. Eight illumination-detection polarization configurations are compared. The tubing was illuminated using 633 nm. Overlay SNR values represent the ratio of Mean Intensity _FEP_ / Mean Intensity _DI_. Scale bar = 100 μm.

### Stage-scanned imaging of iH-MSCs attached to gelMA microcarriers

Flow scanning of cells attached to microcarriers was simulated by embedding fixed iH-MSCs cultured on microcarriers in agarose and loading them into 1 mm I.D. FEP tubing. Raw 1 µm spaced frames acquired on the PLµTo system using 488 nm and 633 nm illumination for CTG and DRAQ-5 fluorescence, respectively, and 633 nm elastic scattering contrast are shown in Figure 5. Although the FEP tubing is visible in each frame, it does not cause significant shadowing artifacts that would degrade illumination and image quality. Cellular morphological features can be seen throughout the CTG volume (Figure 5a). The elastic scattering modality was operated in the left circular polarized illumination and linear vertical polarized detection configuration to minimize the scattering from the FEP tubing while maximizing cell signal (Figure 5b). Using 633 nm illumination and the laser blocking emission filter permits sub-cellular visualization of DRAQ-5-labeled cell nuclei (Figure 5c). Light sheet fluorescence microscopy and tomography provide similar visualization of iH-MSC morphology throughout the microcarrier aggregate (Figure 5d). This aggregate was comprised of 12 microcarriers which were enumerated using the 3D elastic scattering data, and 23 cells that were enumerated using the 3D DRAQ-5 data. The average cell volume was measured to be 4,556 and 3,677 µm^3^ quantified via CTG and elastic scattering, respectively.

**Figure 5:**
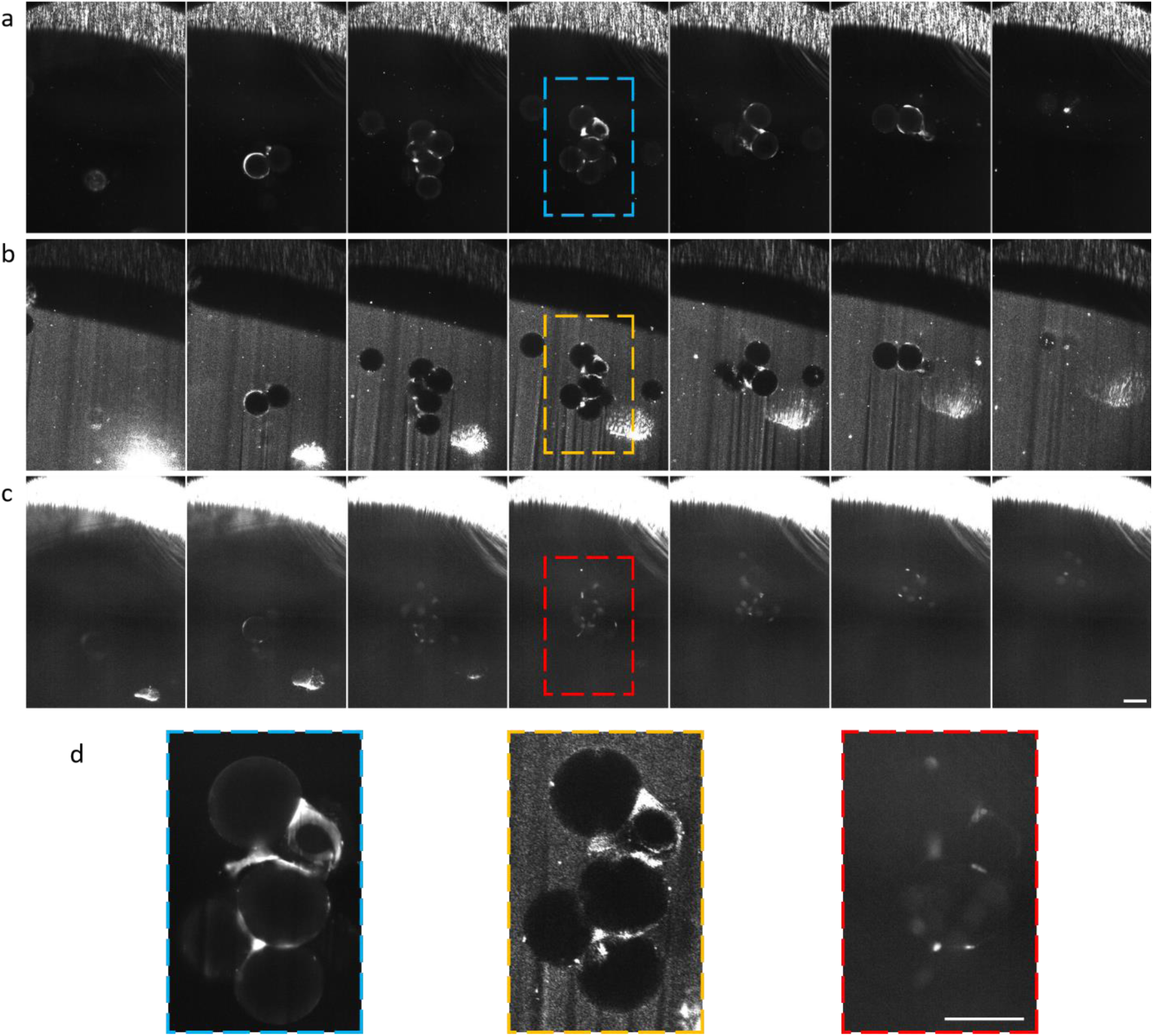
Montage of individual slices of fixed iH-MSCs cultured on gelMa microcarriers and embedded in agarose within 1 mm ID FEP tubing. Slices were acquired with 1 µm spacing and displayed frames here are spaced approximately 27 µm apart through the volume. a) 488 fluorescence of CTG-labeled cells, b) elastic scattering contrast using left circular polarized illumination and linear vertical polarized detection, and c) nuclear-bound DRAQ5 fluorescence imaged with 633 nm illumination. d) Zoomed in ROI showing visualization of cytomorphology and nuclear density within the CTG fluorescence, elastic scattering, and DRAQ5 fluorescence images. Scale bar = 100 µm.

### Application of PLµTo for on-line, non-invasive, and non-destructive imaging of MSCs attached to microcarriers

To demonstrate the applicability of PLµTo as an on-line PAT for monitoring microcarrier-based cell cultures, fixed ihMSCs attached to gelMA microcarriers were pumped with a syringe pump (KDS 100, KD Scientific) at 1,000 nL/min from a 100 mL vertical wheel (VW) bioreactor vessel (PBS-0.1, PBS Biotech) through FEP tubing and imaged at 100 FPS using PLµTo (Figure 6). A montage of images acquired of a populated microcarrier aggregate shows the aggregate coming in and out of the PLµTo image plane (Figure 6b). The left circular illumination and vertical detection polarization states allow for the FEP scattering to be minimized while preserving the cell signal of interest. Highly-scattering cells are discernible on the microcarrier surface and aggregate interior (Figure 6c). The iH-MSC cells can be visualized and segmented from the FEP, microcarrier, and DI water background scattering (Figure 6d).

**Figure 6.**
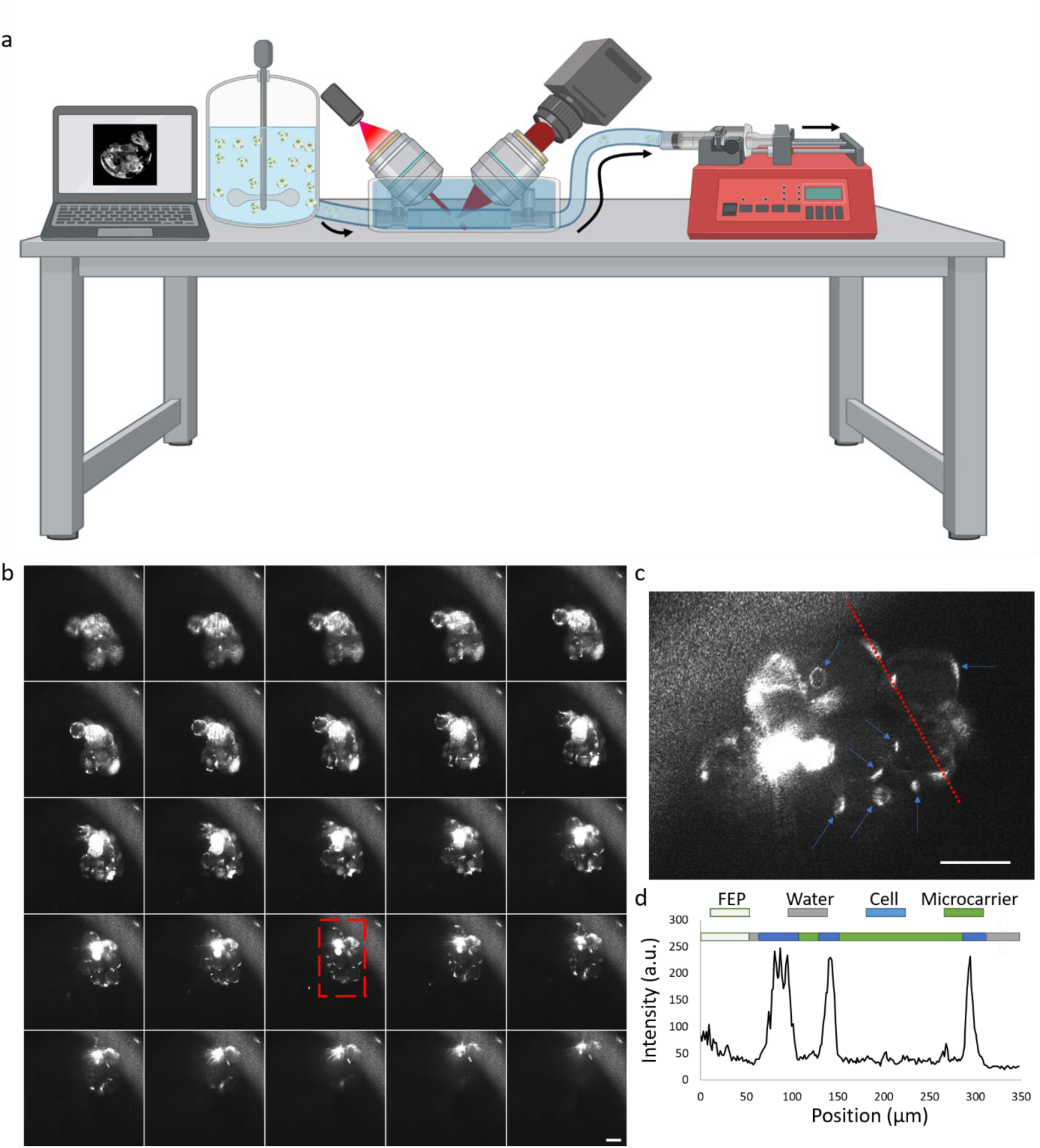
a) Graphic of on-line PLµTo of fixed ihMSCs. b) Montage of cropped regions of interest within scattering contrast images of passage 4 day 3 iH-MSCs attached to 100 µm diameter gelMA microcarriers flowing through FEP tubing. The spheres are illuminated with 633 nm and left circular illumination and vertical detection polarization states were used. The volume flow rate was set to 1000 nL/min, and 100 frames per second were acquired. Frames were acquired with approximately 1 µm spacing but displayed frames are separated by approximately 8 µm along the axial dimension. (c) Zoomed in ROI to visualize cellular structures (blue arrows) on and within the microcarrier aggregate. (d) Intensity versus position projection along the scattering-contract image in (c) showing intensity difference between cells, microcarrier scatter, and background.

## Discussion

The combination of two distinct imaging modalities, fluorescence microscopy for compatibility with standard assays and elastic scattering tomography for label-free contrast assessment of microcarrier-based iH-MSC proliferation, provides a versatile imaging platform for rapid structural and biochemical imaging. The large FOV of the PLµTo system compensates for its moderate resolution; these two specifications are balanced to better enable visualization and quantification of cell number of large microcarrier aggregates *in toto.* The PSF obtained using the PLµTo system are comparable to those obtained by other inverted light-sheet systems; Glaser et al. reported a multi-immersion open-top light-sheet microscope with a 3.5 µm resolution along the collection axis and a hybrid open-top light-sheet microscope with PSFs < 4 µm in each dimension [24]. The measured resolution of the PLµTo system is high enough to allow visualization of sub-cellular morphological features including the nucleus while minimizing the amount of imaging data necessary to image and analyze large microcarrier aggregates.

One advantage of closed-loop in- and on-line PATs is that sterility and yield of the culture process can be maintained using dedicated sampling ports or lines connected to the bioreactor. Here, FEP material, due to its refractive index = 1.34, was used as a sampling line connected to the bottom port of a VW bioreactor and as the cover/window of a microfluidic chip. The FEP tubing and sheet compatibility required characterization to optimize the imaging parameters and conditions for imaging cells under flow in a closed loop system. The incorporation of polarization optics allows for more precise control and optimization of the imaging conditions. By using waveplates in rotation mounts, control over the illumination and detection polarization states allowed for the scattering from the FEP tubing material to be minimized, while preserving cellular signals of interest. This enhances the capabilities of LST by minimizing unwanted background signals (Figures 4-6). The combination of non-invasive LST and image analysis with microfluidics for non-contact sample handling enables a novel PAT for monitoring morphology and density of cells attached to microcarriers.

The FEP material does not fluoresce intensely enough to affect the image quality of the fluorescence modality. Compatibility with standard fluorescence-based assays is valuable when developing novel imaging and analysis methods, and PLµTo is amenable to a wide-range of fluorescence markers that can be illuminated with 488 and 633 nm light. Similarly, FEP produces manageable background scatter that can be minimized using polarization optics. The further development of microfluidics and hydraulic sampling would make this method more robust and reliable. Optimization of the flow mechanisms and flow rates are needed to force the microcarriers to move at controlled rates that do not introduce significant motion blur into the image. It is also possible to fine tune the refractive index (mis)match of the immersion and flow solution in a non-toxic manner to improve the imaging depth and optical resolution [25]. Additionally, the ability to perform multi-scale imaging for microcarriers and aggregates of different sizes would improve the utility of the system as microcarrier aggregates greater than 800 µm in a lateral dimension cannot be imaged fully [9]. An upgraded PLµTo system could incorporate zoom optics for variable magnification and FOVs, pivot scanning for reducing shadow artifacts, and dual-sided illumination for more uniform imaging of large aggregates.

Here, flow-enabled PLµTo was demonstrated as a non-invasive PAT for quantitative monitoring of microcarrier-based stem cell morphology and density. Both contrast modalities, LSFM and LST, are able to visualize 3D objects translated using hydraulic flow. The incorporation of microfluidics and hydraulic flow for volumetric acquisition increases the imaging throughput of the system and creates a hybrid volumetric optical imaging and flow cytometry system. Imaging flow cytometers or volume flow cytometers have been realized in the last decade, but to this point have used fluorescence contrast to image single cells in flow rather than cells attached to microcarriers [26–34]. This demonstration of flow-based PLµTo showed the advantages of moderate resolution and large FOV for imaging and quantification of cells on microcarrier aggregates. This work used label-free elastic scattering contrast and hydraulic flow and is, to the best of our knowledge, the first demonstration of scattering-based volume imaging flow cytometry.

## Conclusion

The work here has culminated in an optical imaging system that can visualize cells attached to microcarriers using fluorescence and elastic scattering contrast and is compatible with microfluidic chips and hydraulic flow for non-destructive and high-throughput imaging. PLµTo can perform in-line fluorescence and elastic scattering assay for monitoring microcarrier cell cultures, and can be utilized to image horizontally-mounted samples. The use of label-free contrast is crucial for non-destructive monitoring as it allows cells to be returned to the bioreactor culture after imaging, preserving the sterility and yield of the culture. The PLµTo system has the capability to image and resolve cytomorphological features of microcarrier cell cultures, and these results demonstrate the PLµTo system as a PAT for on-line monitoring of bioreactor-microcarrier-based cell cultures.

## Author Declarations

Oscar R. Benavides: Conceptualization; Investigation; Writing- original draft; Writing- review & editing.

Berkley P. White: Investigation; Writing- review & editing.

Roland Kaunas: Funding acquisition; Writing- review & editing.

Carl A. Gregory: Funding acquisition; Writing- review & editing.

Alex J. Walsh: Supervision; Funding acquisition; Visualization; Writing- review & editing.

## Data Availability

Data available on request from the authors.

